# Direct selection under resource allocation trade-offs enables the evolution of obligate and facultative sexuality

**DOI:** 10.64898/2026.01.29.702710

**Authors:** Kuangyi Xu

## Abstract

Sexual reproduction is considered to incur multiple costs relative to asexual reproduction. Although previous research has identified indirect selective advantages that help explain the widespread occurrence of sex, the emergence and maintenance of high rates of sexual reproduction remain a central puzzle in evolutionary biology, because indirect selection favoring sex becomes weak when the sexual rate is high. Using a modifier framework that allows the simultaneous evolution of sexual rate and sex allocation, I investigate the evolution of sex via direct selection by incorporating resource allocation trade-offs between sexual and asexual reproduction. The results show that such trade-offs can substantially facilitate the invasion of sex. Crucially, the evolutionarily stable sexual rate depends on the return exponents of female fertility and asexual fertility with respect to resource investment, as well as on the initial allocation to female function within sexual reproduction. Obligate sexuality is evolutionarily stable when asexual fertility exhibits linear or accelerating returns on investment and when both the initial sexual rate and female function allocation within sexual reproduction exceed certain levels. In contrast, facultative sexuality will be evolutionarily stable when female fertility within sexual reproduction and/or asexual fertility exhibit diminishing returns. Contrary to previous theoretical predictions, self-fertilization often inhibits the evolution of sex or reduces the evolutionarily stable sexual rate. This study provides insights into the prevalence of high sexual rates, as well as the continuous spectrum of sexual rates in some groups, highlighting the importance of key parameters in reproductive ecology in shaping the evolution of sex.

**Significance statement:** Why sexual reproduction is so common despite its substantial costs remains a longstanding puzzle in evolutionary biology. Most previous explanations focus on indirect genetic benefits of sex, which generally become weaker as the rate of sexual reproduction increases, and therefore cannot explain the prevalence of intermediate to high sexual rates in nature. This study shows that with resource allocation trade-offs between sexual and asexual reproduction, obligate or facultative sexuality can evolve and be stably maintained via direct selection alone, depending on the marginal returns of asexual and female fertility, as well as the initial allocation to female function. These results highlight the underappreciated role of reproductive ecology in shaping the evolution of sex in nature.

## Introduction

The evolution of sexual reproduction has long attracted interest in evolutionary biology, as sexual reproduction is prevalent despite incurring several costs compared to asexual reproduction (Lehtonen et al. 2012, Meirmans et al. 2012). One major cost of sex is the “cost of males” in separate-sex populations, as females invest resources into production of male individuals, which invest minimally into sired offspring (Smith & Maynard-Smith 1978). Another major cost is the “cost of meiosis” (Williams 1975, Lewis 1987), as asexuals transmit the entire genome to offspring, while sexuals transmit only half of the genome to their offspring. These costs have been reported in various taxa (Michaels & Bazzaz 1986, Innes et al. 2000, Wolinska & Lively 2008). There are other additional costs of sex, including the cost related to sexual selection, mate limitation and sexually transmitted diseases (Meirmans et al. 2012). Moreover, within populations, sexually reproduced offspring tends to be less fit than asexually reproduced individuals due to recombination and segregation (Charlesworth & Barton 1996, Otto 2003, Xu 2024). Therefore, the invasion of sex requires sexually produced offspring to be fitter than asexually produced offspring to offset the cost (Maynard Smith 1978; Charlesworth 1980; Bulmer 1982), but models considering beneficial alleles or deleterious mutations often predict the opposite (Peck & Waxman 1999; Otto 2003; Xu 2024). Although factors such as self-fertilization and a high female-to-male offspring ratio can mitigate the costs of sex, these factors are generally insufficient to reverse the direction of selection against sex (Charlesworth 1980; Uyenoyama 1984; Joshi & Moody 1995, 1998; Xu 2024).

To reconcile this contradiction, studies in the past decades have focused on identifying selective advantages of sex over asexual reproduction via segregation and recombination (reviewed in Otto 2009; Hartfield & Keightley 2012; Otto 2021). One category of models considers in infinitely large populations and characterizes conditions in which selection depletes genetic variation and mean fitness increases with greater genetic variation, allowing sex to invade by increasing genetic variation (Barton 1995; Otto 2003). A second category of models shows that sex can be favored when selection varies over time or space (Gandon & Otto 2007; Becks & Agrawal 2010; Lively 2010). Nevertheless, these deterministic models suggest that sex is generally favored only under restricted conditions (Otto & Michalakis 1998; Otto 2009). Therefore, a third category of models considers finite populations and focuses on the benefits of sex from reducing selective interference, where selection at one locus is rendered ineffective by selection at linked loci due to interactions between selection and genetic drift. By bringing together multiple beneficial alleles onto a haplotype, sex can spread along with favorable genotypes (Barton & Otto 2005; Roze & Michod 2010), and the evolution of sex via selective interference is considered a promising solution, as it occurs across a wide range of conditions (Otto 2021). Other selective advantages of sex can occur via sexual selection (Agrawal 2001) and sexual conflict (Burke & Bonduriansky 2019).

While the above studies can explain the occurrence of sex, they are ineffective in explaining why sex is the dominant mode of reproduction in many lineages of multicellular eukaryotes, which is considered the “true paradox of sex” (Green & Noakes 1995; Burke & Bonduriansky 2017; Hörandl 2024), nor the continuous spectrum of intermediate sexual rates in many primitive eukaryotes, as well as in some animals and plants (Bell 1982, Aliyu et al. 2010). In fact, previous studies show that a mixture of sexual and asexual reproduction (i.e., facultative sexuality) cannot be stably maintained by direct selection (Charlesworth 1980; Roughgarden 1991; Joshi & Moody 1995); even when indirect selection is incorporated, a very low rate of sex tends to be optimal or evolutionarily stable, balancing the direct costs and indirect benefits of sex (Charlesworth et al. 1993; Green & Noakes 1995; D’Souza & Michiels 2010; Hojsgaard & Hörandl 2015; Kokko 2020; Stetsenko & Roze 2022). One frequently invoked explanation for the maintenance of obligate sexuality is the physiological constraint on reverting to asexual reproduction, which can be costly once sex is established (Corley & Moore 1999; Engelstädter 2008). However, this hypothesis does not explain how obligate sexuality evolved in the first place, the frequent transitions between obligate sexuality and predominant asexuality (Schwander et al. 2011; Neiman et al. 2014), or the maintenance of facultative sexuality in nature.

Thus, the answer to the maintenance of high sexuality, as well as the continuous spectrum of sexual rates, may still lie in understanding direct selection on sex. One factor that may be crucial in shaping direct selection on sex, but remains not well understood, is the trade-off in resource allocation (Charnov 1982). Most previous models implicitly assume that increased asexuality does not affect male gamete production. Nevertheless, in reality, individuals exhibiting higher levels of asexuality are often associated with reduced sexual function (Sutherland et al. 1988; Mogie 1992; Jarne & Auld 2006; Whitton et al. 2008; Schwander et al. 2011; Stelzer & Lehtonen 2016). In fact, models show that the cost of sex is substantially lower when asexuals have reduced male gamete output (Joshi & Moody 1995, 1998). More generally, the evolution of alleles that modify resource allocation between asexual and sexual reproduction should depend on how allocation to female and male functions is altered.

Here, using a modifier model, I investigate how trade-offs in resource allocation shape the evolution of sex and determine evolutionarily stable sexual rates. The model considers a diploid population with a sex modifier locus carrying two alleles, *A* and *a*. Individuals allocate fixed reproductive resources to sexual and asexual reproduction (Figure 1a). For genotype *AA*, the proportions allocated to sexual and asexual reproduction are *σ* and *σ*_*a*_ = 1 − *σ*, respectively, and within sexual reproduction, resources are split between female and male functions as *σ*_*f*_ = *Fσ* and *σ*_*m*_ = (1 − *F*)*σ*. These resources include both gamete production and structures or behaviors necessary for sexual function (e.g., flower displays). For genotypes *Aa* and *aa*, allocations to sex are *σ* + *h*Δ*σ* and *σ* + Δ*σ*, respectively, where Δ*σ* represents the modification of sexuality and *h* is the dominance coefficient. Within Δ*σ*, the proportions devoted to female and male functions are *f* and 1 − *f*. When *f* ≠ *F*, modification in sexual rate is accompanied by changes in sex allocation. Resource allocations for all three genotypes are summarized in Figure 1c.

**Figure 1.**
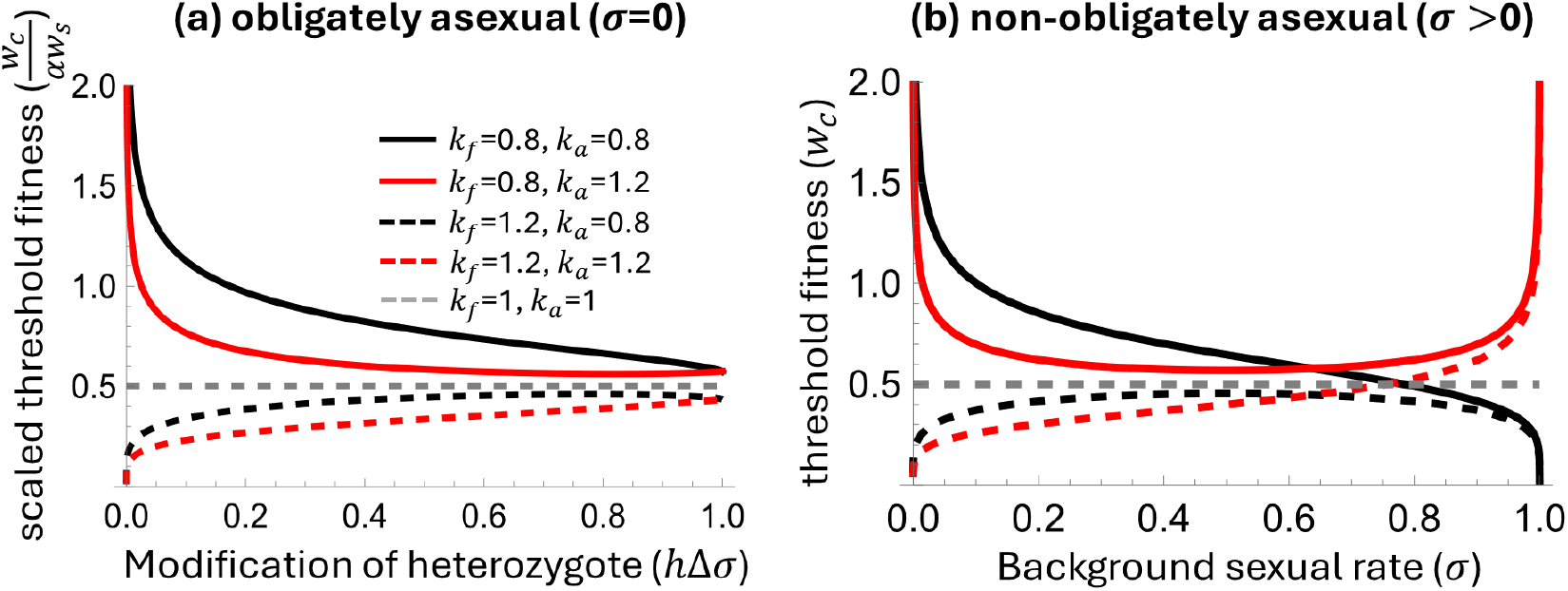
Invasion condition of sex in an obligately asexual population (panel a) and non-obligately asexual population (panel b) under different return exponents of female fertility and asexual fertility. Sex can invade when the scaled fitness of asexual offspring is below a threshold fitness *w*_*c*_, which is scaled by the rate of viable selfed offspring *αw*_*s*_ in panel (a). For both cases, the likelihood of invasion of sex depends on whether female fertility, *k*_*f*_, is below 1 or not. The invasion condition also depends on the modification magnitude (x axis in panel (a)) and the background sexual rate (x axis in panel (b)). Results are model predictions from Equations (1) and (2) with *f* = *F* =0.5 and *α*=0 for panel (b).

Fertility via a given reproductive mode increases with resource allocation and also depends on the ecological context of reproduction. I define female fertility as the number of female gametes (in hermaphroditic populations) or sexual female offspring (in separate-sex populations) of an individual that are successfully fertilized. Male fertility refers to the number of male gametes successfully dispersed or the number of male offspring that successfully mate. I assume that asexual fertility, female fertility, and male fertility are power functions of resource allocation as 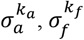, and 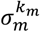, respectively (Figure 1a) (Charnov 1979; Charlesworth & Charlesworth 1981; Dorken et al. 2025). Here the return exponent 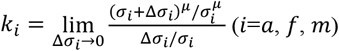 is the percentage increase in fertility with a percentage increase in resource investment. When the return exponent is lower, fertility is higher at low investment but saturates more quickly as resource allocation increases (Figure 1b). Since investment in female fertility in the current model includes both resources for gamete production and those required for successful fertilization, *k*_*f*_ may differ from *k*_*a*_, which can occur if sexual and asexual reproduction occurs through different developmental processes (e.g., seed versus vegetative reproduction). Also, *k*_*a*_ and *k*_*f*_ may deviate from linearity. For example, female fertility may exhibit diminishing returns in plant populations under pollen limitation (Knight et al. 2005) or when ovule fertilization rates increase deceleratingly with flower display size (Grindeland et al. 2005; Karron & Mitchell 2012), whereas accelerating returns may occur if there is competition among females for mates (Johnstone et al. 1996). For male fertility, plant populations are generally expected to show diminishing returns (*k*_*m*_ < 1) (Dorken et al. 2025; Harder & Johnson 2025), whereas in animals, male fertility may exhibit accelerating returns if there is competition for siring success. In previous models, the returns of asexual and female fertility are often assumed to be linear (*k*_*a*_ = *k*_*f*_ = 1; Charnov 1979; Joshi & Moody 1995), and as will be shown, this assumption critically influences the results.

Let *r*_*a*_ be the resource cost of an asexually produced offspring relative to a sexually produced offspring. Thus, the number of asexual offspring is 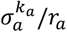, and the number of sexual offspring via the female and male functions is 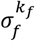 and 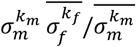, respectively, where 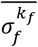 and 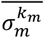 are the population averages. Under isogamy, *r*_*a*_ = 2, and under complete anisogamy, *r*_*a*_ = 1, and other values are possible when sexual and asexual reproduction occur via different mechanisms (e.g., seed vs. vegetative reproduction). To incorporate self-fertilization in hermaphrodites, a fraction *α* of female gametes are assumed to be self-fertilized, while 1 − *α* are outcrossed, and the proportion of male gametes used for selfing is negligible. To account for fitness differences among offspring from different modes (Xu 2024), the fitness of asexually produced and self-fertil ized offspring, relative to outcrossed offspring, is denoted *w*_*a*_ and *w*_*s*_, respectively. In the result section, I describe results assuming a hermaphroditic population, since the outcomes are identical for both hermaphroditic and separate-sex populations (see *Materials and Methods*).

## Results

### Invasion conditions

A sex-enhancing modifier (Δ*σ* > 0) invades when the scaled relative fitness of asexual offspring, 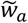, is below a threshold *w*_*c*_, where 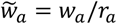 captures differences in both fitness and resource cost to reduce parameters. The invasion condition of sex differs depending on whether the residential population is obligately asexual or not. For intuition, when residents are obligately asexual, rare sexually reproducing mutants cannot have their male gametes fertilize others, and their female gametes cannot be fertilized unless via self-fertilization.

When the original population is obligately asexual (*σ* = 0), the invasion of sex requires (Section 1 of SI Appendix)

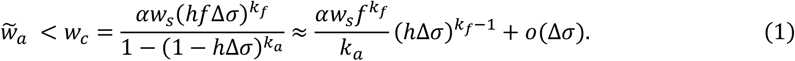

In the middle expression, the denominator represents the fitness loss of the mutant due to reduced asexual reproduction, while the numerator represents the fitness gained through self-fertilization. The likelihood of sex invading an obligately asexual population depends critically on the return exponent of female fertility, *k*_*f*_, as well as the magnitude of sexual rate modification, Δ*σ*. Invasion is more likely when female fertility exhibits diminishing returns with investment (*k*_*f*_ < 1) and the modification effect is small (solid line, Figure 1). In contrast, invasion is generally much more difficult when female fertility shows linear or accelerating returns with investment (*k*_*f*_ ≥ 1; dashed line, Figure 1).

When the residential population is facultatively sexual (*σ* > 0), the invasion condition of sex is (Section 1 of SI Appendix)

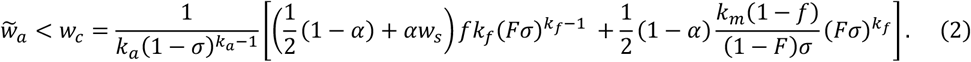

The term 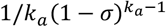 represents the cost from reduced asexual reproduction. he two terms in the square brackets represent the fitness benefits to the mutant from increased female and male fertility per unit of investment in sexual reproduction, respectively, with the coefficient 1/2 accounting for the cost of meiosis during outcross fertilization. Equation (2) clarifies that the classical “twofold cost of sex” (i.e., *w*_*a*_ /*r*_*a*_ < 0.5) hold only under very specific conditions, which requires equal allocation to female and male functions (*F* = 0.5), linear returns in fertility for all reproductive modes (*k*_*a*_ = *k*_*f*_ = *k*_*m*_ = 1), no self-fertilization (*α* = 0), and complete anisogamy (*r*_*a*_ = 1).

The background sexual rate *σ* affects both the cost from reduced asexual fertility and the benefit from increased female and male fertility, and therefore the invasion likelihood sex (Figure 1b), with the impacts of *σ* depending critically on the return exponent of female fertility *k*_*f*_ and asexual fertility *k*_*a*_ (see Section 2 of SI Appendix). In general, sex can invade readily when *σ* is low and female fertility exhibits diminishing returns (low *k*_*f*_; solid lines in Figure 1b), or when *σ* is high and asexual fertility exhibits accelerating returns (high *k*_*a*_; red lines for *σ* > 0.8 in Figure 1b). Under linear returns (*k*_*a*_ = *k*_*f*_ = 1), both the cost and benefit, and hence the invasion condition, are independent of the background sexual rate (gray dashed line, Figure 1b). When *k*_*a*_, *k*_*f*_ < 1, a higher *σ* inhibits the invasion of sex because it reduces the benefit from increased female and male fertility while strengthening the cost (black solid line, Figure 1b). Conversely, when *k*_*a*_, *k*_*f*_ > 1, a higher background sexual rate promotes invasion by increasing the benefit and reducing the cost (red dashed line, Figure 1b). When *k*_*f*_ > 1 > *k*_*a*_ or *k*_*f*_ < 1 < *k*_*a*_, increasing *σ* simultaneously affects both cost and benefit, producing a non-monotonic effect on invasion. Specifically, when *k*_*f*_ < 1 ≤ *k*_*a*_, increasing *σ* from 0 initially inhibits invasion and then promotes it when *σ* > (1 − *k*_*f*_)/(*k*_*a*_ − *k*_*f*_) (red solid line, Figure 1b). When *k*_*f*_ ≥ 1 > *k*_*a*_, increasing *σ* from 0 first promotes invasion and then inhibits it when *σ* > (1 − *k*_*f*_)/(*k*_*a*_ − *k*_*f*_) (black dashed line, Figure 1b).

The dependence of sex invasion likelihood on the background sexual rate *σ* is critical for determining whether facultative sexuality can represent an evolutionarily stable state (ESS), as summarized in Table 2 and discussed in the next subsection. When invasion becomes monotonically more likely with increasing *σ* or is independent of *σ*, which occurs when *k*_*a*_, *k*_*f*_ ≥ 1, the evolutionarily stable sexual rate is either obligately asexual (*σ* = 0) or obligately sexual (*σ* = 1). Intermediate sexual rates can potentially be ESS only when higher background sexual rates inhibit the invasion of sex.

The invasion of sex also depends on additional factors (see Section 2 of the SI Appendix). In general, sex is more likely to invade when the return exponent of male fertility *k*_*m*_ is higher (compare solid and dashed lines in Figure 2) and when the background allocation to female function *F* is larger, although the effect of *F* can be non-monotonic when modifications in sexuality increase relative allocation to female function within sexual reproduction (i.e., *f* > *F*; compare red and blue lines in Figure 2). In these cases, the benefit of sex from increased siring success is greater. Moreover, when modifications in sexuality are accompanied by changes in sex allocation, diverting more resources to the male function (lower *f*) promotes invasion when *F* exceeds a critical level, 1/[1 + ((1 − *α*)*k*_*m*_/*k*_*f*_)/(1 + (1 − 2*w*_*s*_)*α*)] (compare solid lines for *F* > 0.5 and dashed lines for *F* > 0.4 in Figure 2), whereas diverting more to the female function (higher *f*) promotes invasion when *F* is below this threshold.

**Figure 2.**
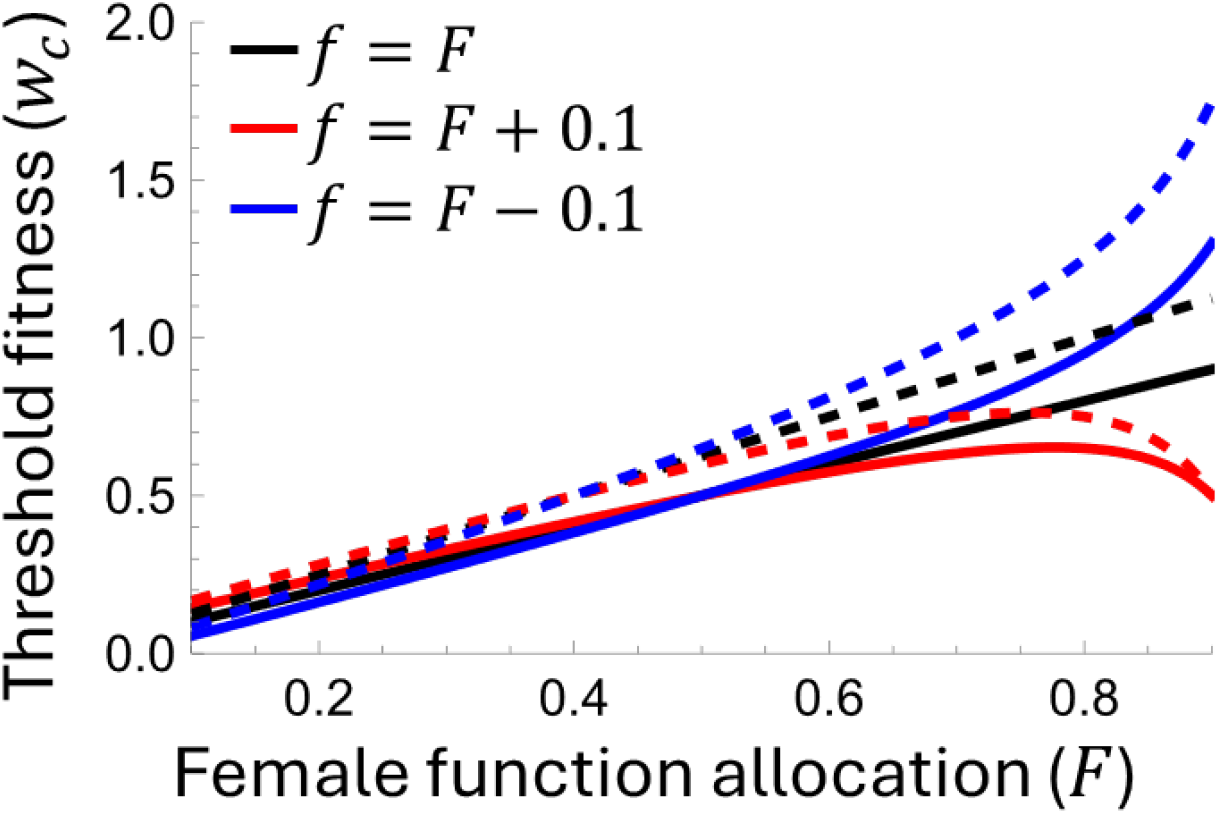
Impacts of the background female function allocation within sexual reproduction (*F*) on the threshold fitness *w*_*c*_. Sex can invade when the scaled fitness of asexually reproduced offspring 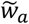 is below *w*_*c*_. Black, red, and blue lines correspond to different degrees of modification to female function *f*, where *f* = *F* means modification of sexuality does not affect sex allocation. Solid and dashed lines show results under different return exponents of male fertility with *k*_*m*_=1 and 1.5, respectively. Results are obtained from Equation (2) with *k*_*f*_ = *k*_*a*_ = 1, *σ* = 0.5, *α* = 0.

### Evolutionarily stable allocation

As noted above, the evolutionarily stable sexual rate depends critically on the return exponents of asexual and female fertility *k*_*a*_ and *k*_*f*_, as they determine whether the feedback between the invasion likelihood of sex and the background sexual rate is positive or negative (Table 2). Below, we discuss four cases depending on whether *k*_*a*_ and *k*_*f*_ are greater than 1. The evolutionary outcome may also depend on the population’s initial state and on the magnitude of the changes introduced by sex-modifier mutants. In general, facultative sexuality can be an ESS when female fertility exhibits diminishing returns (*k*_*f*_ < 1), whereas obligate sexuality may be an ESS when the initial sexual rate and allocation to female function are sufficiently high.

When *k*_*a*_, *k*_*f*_ ≥ 1, populations will evolve to either obligate asexuality or obligate sexuality, depending on the initial level of allocation to female function within sexual reproduction, *F* (case (1) in Table 2). Specifically, a population with initial sexual rate *σ* and female allocation *F* evolves to obligate asexuality when *F* < *F*^∗^(*σ*) (population 1 in panel (1a) of Table 2), and to obligate sexuality when *F* > *F*^∗^(*σ*) (population 2 in panel (1a) of Table 2), where the threshold *F*^∗^(*σ*) is (derived in Section 3 of the SI Appendix)

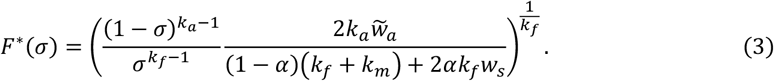

By Equation (3), evolution to obligate sexuality occurs across a larger parameter space (i.e., *F*^∗^ is lower) when *k*_*a*_ is higher and *k*_*f*_ is lower (see panel (1b) in Table 2), and also when the return exponent of male fertility, *k*_*m*_, is higher.

When *k*_*f*_ ≥ 1 > *k*_*a*_, the ESS depends on whether the exponent of asexual fertility, *k*_*a*_, lies above or below a threshold value (indicated by the x-axis position of the dashed line in panel (2c) of Table 2). When *k*_*a*_ exceeds this threshold, populations always evolve toward obligate asexuality (panel (2a) of Table 2), although populations with a high initial allocation to female function may transiently evolve higher sexual rates (population 1 in panel (2a) of Table 2). When *k*_*a*_ is below this threshold, populations evolve toward either facultative sexuality or obligate asexuality, depending on whether the initial allocation to female function *F*is above or below the threshold *F*^∗^given by Equation (3) (panel (2b) of Table 2). Notably, this critical threshold of *k*_*a*_ is generally far below 1 (panel (2c) of Table 2). Therefore, given that the return exponents of asexual and female fertility are expected not to deviate far from 1 in nature, obligate asexuality may often be the ESS in this scenario.

Interestingly, populations with the same initial state may ultimately evolve toward different ESSs, depending on the effect size of modifier mutants. This divergence occurs when the initial allocation to female function *F* lies slightly below the minimum of the critical threshold *F*^∗^ (blue line in panel (2b) of Table 2). When modifier mutants have small effects, changes in *F* are slight, and *F* remains below *F*^∗^as sexuality decreases, leading populations to evolve toward obligate asexuality (red trajectory of population 3 in panel (2b) of Table 2). In contrast, when modifier mutants have large effects, the evolutionary trajectory may cross above the critical threshold *F*^∗^, causing populations to evolve toward facultative sexuality (gray trajectory of population 3 in panel (2b) of Table 2).

When female fertility exhibits diminishing returns with investment (*k*_*f*_ < 1), populations will evolve toward either obligate or facultative sexuality, and obligate asexuality can no longer be an ESS. When *k*_*f*_ < 1and *k*_*a*_ ≥ 1, populations evolve to obligate sexuality provided that the initial sexual rate and allocation to female function are sufficiently high (population 3 in panel (3a) of Table 2); otherwise, they evolve toward facultative sexuality (populations 1 and 2 in panel (3a) of Table 2). In general, the parameter space leading to obligate sexuality expands as the return exponent of asexual fertility (*k*_*a*_) increases (compare black and red lines in panel (3b) of Table 2) and as the return exponent of female fertility decreases (compare solid and dashed lines in panel (3b) of Table 2). For the case when *k*_*f*_, *k*_*a*_ < 1, facultative sexuality is always the ESS, regardless of the initial state (case (4) of Table 2).

When facultative sexuality is ESS, the sexual rate, *σ*^∗^, is given by (derived in Section 2 of SI Appendix)

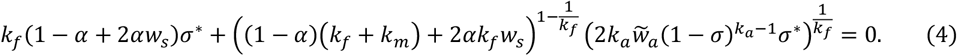

This expression shows that the ESS sexual rate is higher when the return exponent of male fertility *k*_*m*_ is higher, when the return exponents of female and asexual fertility are lower (compare solid lines in panel (2c), round versus square dots in panel (3b), and lines in panel (4b) of Table 2), and when the fitness of asexually produced offspring *w*_*a*_ is lower. For the special case when *k*_*a*_ = 1, the ESS state is

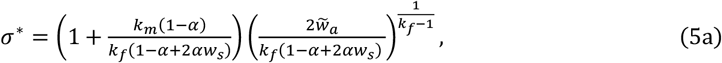

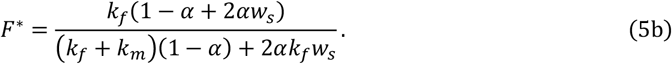

In this case, the allocation to female function at the ESS is independent of the fitness of asexually produced offspring 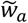 and increases with a larger return exponent of female fertility *k*_*f*_ and a smaller return exponent of male fertility *k*_*m*_. In particular, under random mating, *F*^∗^ = *k*_*f*_/(*k*_*f*_ + *k*_*m*_), which recovers the classical result for ESS sex allocation (Charlesworth & Charlesworth 1981; Dorken et al. 2025).

### Effects of self-fertilization

The possibility of self-fertilization in hermaphrodites can strongly influence both the invasion conditions and the ESS. We have shown in Equation (1) that sex can invade an obligately asexual population only if self-fertilization is possible. When resident populations are not obligately asexual (*σ* > 0), selfing has opposing effects on the evolution of sex: it reduces the cost of meiosis but lowers the fitness of sexually produced offspring due to inbreeding depression (Charlesworth & Willis 2009; Xu 2024). A higher selfing rate promotes the invasion of sex (∂*w*_*c*_/ ∂*α* > 0) when the fitness of selfed offspring is sufficiently high, as given by

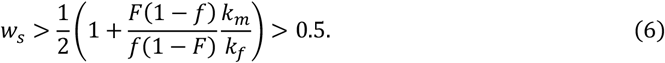

Equation (6) reduces to *w*_*s*_ > (1 + *k*_*m*_/*k*_*f*_)/2 if modifications of the sexual rate nearly preserve the original level of sex allocation (*f* ≈ *F*). Because selfed offspring are typically less fit than outcrossed offspring, selfing can only promote the invasion of sex if male fertility exhibits lower returns than female fertility with resource investment (i.e., *k*_*m*_ < *k*_*f*_), which may be the case in many plant populations (Zhang 2006). Selfing also affects the ESS sexual rate. In general, only when the relative fitness of selfed offspring is sufficiently high does selfing increase the likelihood that populations evolve toward facultative or obligate sexuality and potentially increase the ESS sexual rate. Otherwise, selfing favors obligate asexuality as the ESS and reduces the ESS sexual rate (see Section 4 of the SI Appendix).

## Discussion

This study integrates resource allocation trade-offs between asexual and sexual reproduction and non-linear relationships between fertility and resource investment into a modifier framework that allows the evolution of both sexual rate and sex allocation. With resource allocation trade-offs, direct selection alone may allow sex to invade readily, as well as the evolution and maintenance of obligate or facultative sexuality across a wide range of conditions. The results highlight that key parameters in reproductive ecology are critical in determining the evolutionary outcomes of sexual rates, as summarized in Table 2. Obligate sexuality may be evolutionarily stable when asexual fertility exhibits linear or accelerating returns (*k*_*a*_ ≥ 1), while obligate asexuality may be evolutionarily stable when female fertility exhibits linear or accelerating returns with resource investment (*k*_*f*_ ≥ 1), and facultative sexuality may often be ESS when female fertility exhibit diminishing returns (*k*_*f*_ < 1). The key intuition is that the return exponents of female and asexual fertility determine whether the invasion likelihood of sex increases or decreases with the background sexual rate. These results contrast with previous conclusions that the direct cost of sex is generally prohibitive and that facultative sexuality cannot be evolutionarily stable (Maynard Smith 1978; Charlesworth 1980; Bulmer 1982; Joshi & Moody 1995).

The results identify several conditions that promote the evolution towards obligate sexuality or high sexual rates. First, high allocation to the female function relative to the male function within sexual reproduction, which extends the findings from previous studies on the role of sex ratio (Charlesworth 1980; Joshi & Moody 1995, 1998). Second, female fertility exhibits substantial diminishing return with investment. Third, high return exponent of male fertility with resource investment. Accelerating returns may occur if insufficient resource investment leads to disproportional reduction in the viability or mating success of male gametes or offspring, which may likely occur under intrasexual and intersexual selection (Iwasa & Pomiankowski 1991; Doorn & Weissing 2006; Jaquiéry et al. 2010; Georgiev et al. 2015), which is an additional mechanism for sexual selection to mitigate the cost of sex (Agrawal 2001; Siller 2001; Kleiman & Hadany 2015). However, plant populations are generally expected to exhibit diminishing returns in allocation to male function (Dorken 2025, Harder & Johnson 2025), which may partly explain why asexual reproduction is more common in plants than in animals.

Many plant and animal clades exhibit frequent transitions between obligately sexual and predominantly asexual lineages (Neiman et al. 2014). The current results suggest these shifts may be triggered by initial perturbations in the relative allocation between female versus male function, or by changes in the return exponents of female and male fertility. Alternatively, transitions from obligate sexuality to predominant asexuality can also occur if modifier mutations inducing asexuality leads to a substantial reduction in female function.

Moreover, the current results suggest that animal and plant groups maintaining intermediate sexual rates (Bell, 1982; Aliyu et al., 2012) may experience diminishing returns of female fertility (*k*_*f*_ < 1), and that variation in their sexual rates among populations may be partly driven by differences in the return exponents of fertility associated with different reproductive modes. Importantly, we show that intermediate sexual rates can be evolutionarily stable under direct selection alone, in contrast to previous predictions (Charlesworth, 1980; Joshi & Moody, 1995). This discrepancy arises because earlier models typically assumed linear returns of asexual and female fertility with resource investment and restricted modifier alleles to producing either obligate sexual or obligate asexual reproduction, rather than allowing continuous modification of sexual rates. Such assumptions precluded the detection of feedback between direct selection on sex and background sexual rates. It should be noted, however, that additional factors may influence the evolution of facultative sexuality and the stable sexual rates, including sexual antagonism, mate limitation, dispersal, and genetic drift (Burke & Bonduriansky, 2017, 2019; Tanaka & Bonduriansky, 2025).

In contrast to previous finding that selfing tends to mitigate the cost of sex (Charlesworth 1980, Uyenoyama 1984), incorporating resource trade-offs in the current model suggests that selfing may often inhibit both the invasion of sex and makes the evolution towards obligate sexuality less likely, unless the fitness of selfed offspring relative to outcrossed offspring is sufficiently high (at least 0.5). Regarding the expectation that higher selfing rates can purge inbreeding depression (Lande and Schemske 1985; Roze 2015), low to intermediate selfing rates may inhibit the evolution of sex via direct selection, whereas high selfing rates may potentially promote it. Interestingly, this result coincides with the finding that the indirect selective advantage of sex is reduced in populations with intermediate selfing rates but greatly increased in highly selfing populations (Xu 2024). The results appear to be partly inconsistent with the observed negative association between selfing and asexual reproduction (Barrett 2002; Vallejo‐Marín et al. 2007; Whitton et al. 2008). This may be because selfing is often accompanied by changes in a suite of correlated traits, known as the “selfing syndrome” (Sicard & Lenhard 2011), and these associated changes often reduce the cost of sex (e.g., selfers tend to have smaller floral displays). In addition, selfing can provide reproductive assurance under mate or pollinator limitation, and selfers typically allocate more resources to female function than outcrossers (Ritland & Ritland 1989; Parachnowitsch & Elle 2004), which inhibits the evolution of asexual reproduction.

Several factors are not explicitly incorporated in the current model. First, the model assumes identical resource allocation among individuals, whereas reproductive resources may vary between individuals due to spatial heterogeneity (Sugiyama and Bazzaz 1998; Lim et al. 2014) or trade-offs between growth and reproduction (Doust 1989; Heino and Kaitala 1999). Previous studies on condition-dependent sex show that the evolution of sex is promoted when individuals in poorer condition allocate a greater fraction of resources to sexual reproduction (Redfield 1988; Gessler and Xu 2000; Hadany and Beker 2003; Hadany and Otto 2007). In addition, the model does not explicitly account for the selective disadvantage of sex arising from mate limitation (Xu 2024). However, the effect of mate limitation may be effectively equivalent to reduced offspring fitness or fertility, and can therefore be incorporated through the parameter 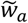. Finally, it should be noted that the current predictions are based solely on direct selection. If indirect selection on sex is also considered, obligate asexuality may not be an ESS, and the ESS sexual rate may be higher.

## Materials and Methods

For hermaphroditic populations, when the population is not obligately asexual (*σ* > 0), the recursions of genotype frequencies *p*_*i*_ (*i* = *AA, Aa, aa*) are

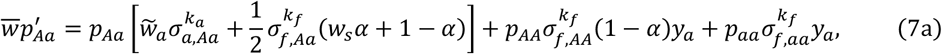

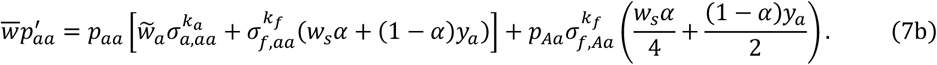

Here 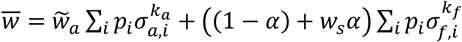 is the mean fitness, and 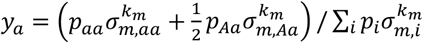 is the frequency of allele *a* among male gametes that successfully fertilize female gametes, with *σ*_*a,i*_, *σ*_*f,i*_, and *σ*_*m,i*_ being the resource allocation to asexual reproduction, female function and male function for genotype *i*, summarized in Table 1(c). The invasion condition of a modifier is determined by the leading eigenvalue of Equations (7a) and (7b) (Section 1 of SI Appendix). If the resident population is obligately asexual (*σ* = 0), Equations (7a) and (7b) still apply, with terms involving 1 − *α* removed, since outcross fertilization is not possible.

**Table 1.**
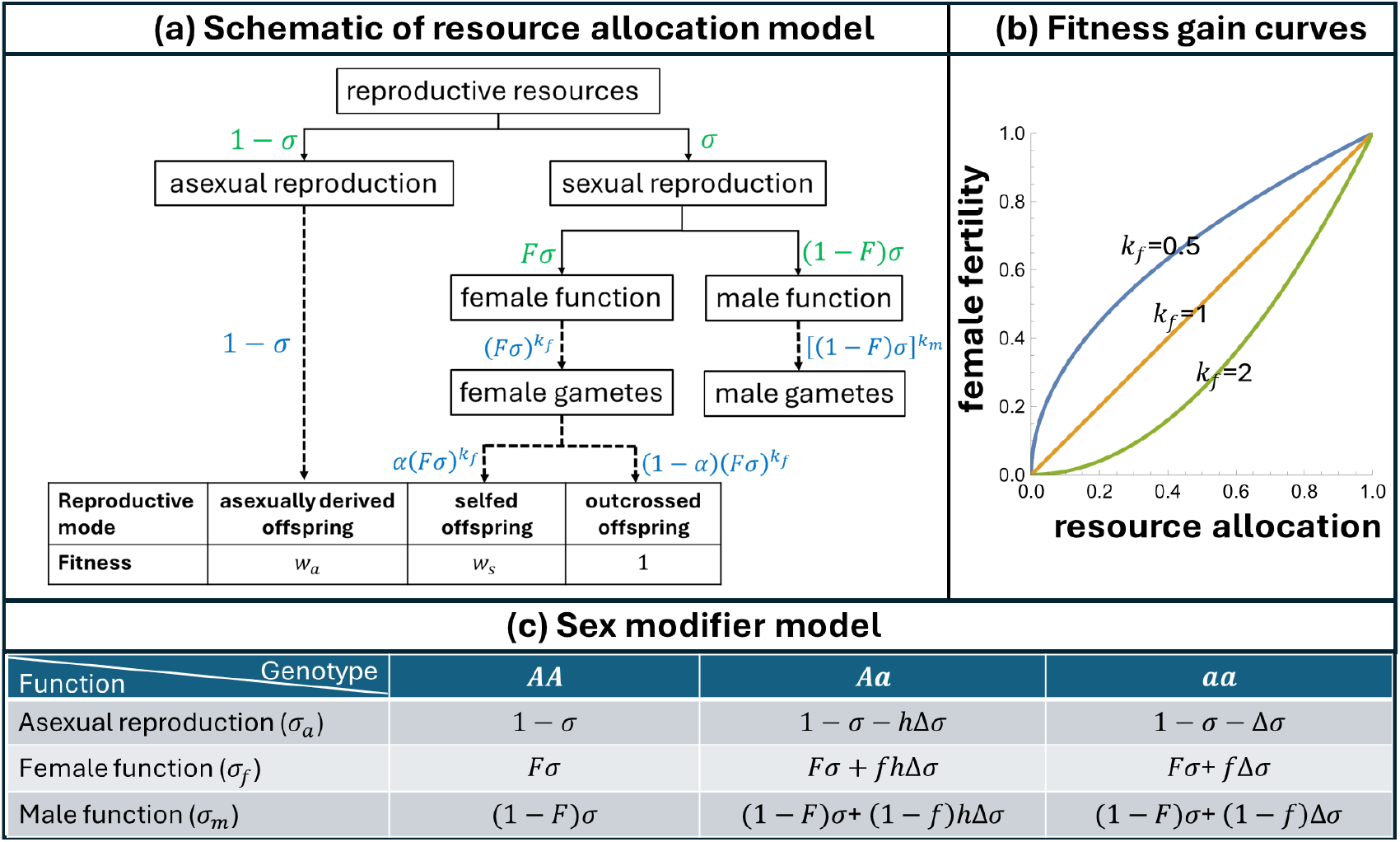
Schematic summary of the sex modifier model with trade-off in resource allocation to different reproductive modes.

**Table 2.**
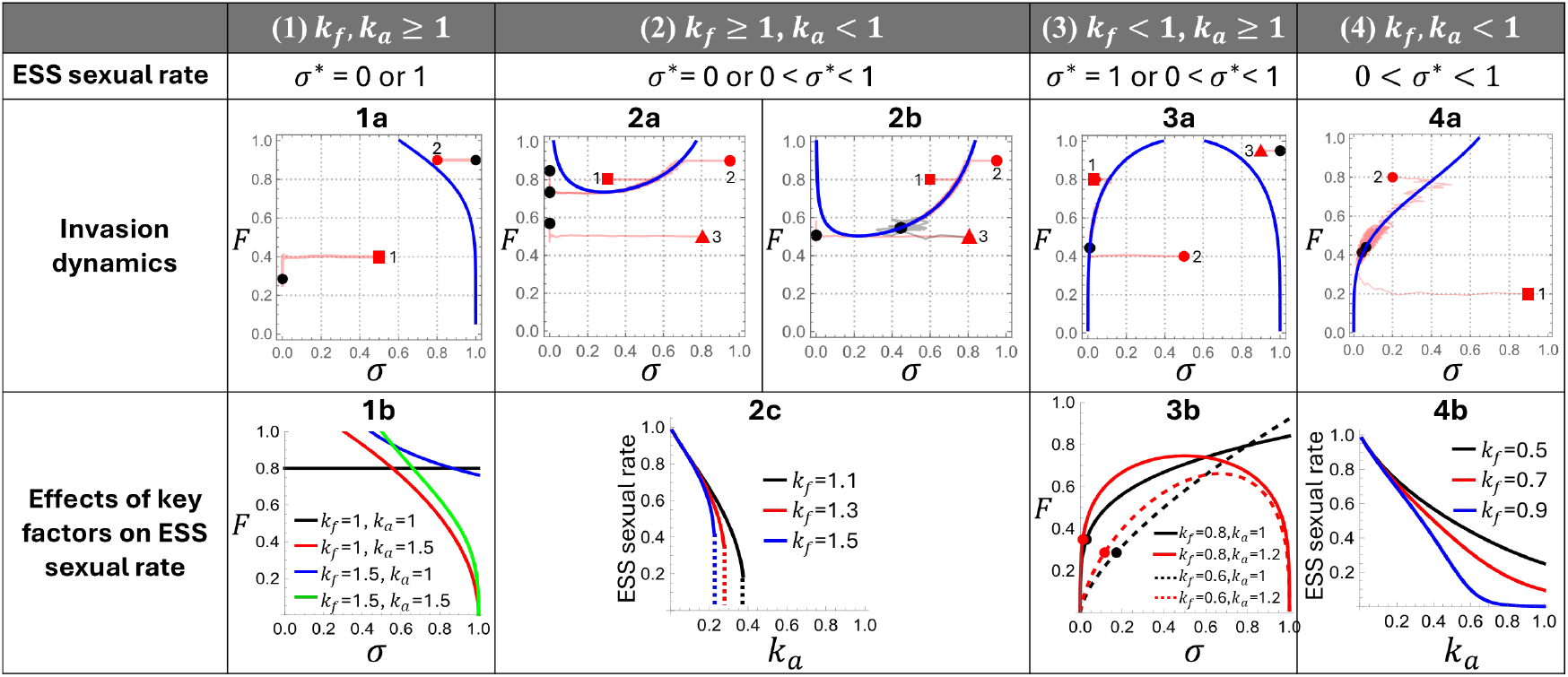
Evolutionarily stable sexual rate and invasion dynamics under 4 cases of return exponents of female fertility (*k*_*f*_) and asexual fertility (*k*_*a*_). In the invasion dynamics panels, blue curves show the critical *F*^∗^ given by Equation (3), above (below) which populations evolve toward higher (lower) sexual rates. Red points with different numbers represent populations with different initial allocation states, and black dots indicate the final states after 20,000 generations. Trajectories are nearly horizontal because the changes in female function allocation *F* occurs via modifications of the sexual rate, which are of second-order effect, *O*(Δ*σ*(*f* − *F*)). The maximum relative magnitude of modification is Δ*σ*_max_ = Δ*F*_max_ = 0.1, except for the gray trajectory in panel (2b), where Δ*σ*_max_ = Δ*F*_max_ = 0.5 (see *Materials and Methods*). The bottom row shows the effects of key factors on the ESS. Panels (1b) and (3b) show the critical *F*^∗^(*σ*) as a function of background sexual rate *σ*. In panel (1b), populations evolve to obligate sexuality when starting above the curve, and to obligate asexuality otherwise. In panel (3c), populations evolve to obligate sexuality when starting with *F* above the maximum of the curve or in the upper-right region of the bell-shaped curve (red lines); otherwise, they evolve to facultative sexuality (dots). In panel (2c), the ESS is obligate asexuality when *k*_*a*_ exceeds the value marked by the dashed line, and the solid lines show the dependency of the ESS sexual rate on *k*_*a*_ when facultative sexuality is the ESS. In panel (4b), facultative sexuality is always the ESS, and the lines show the effects of *k*_*a*_ and *k*_*f*_on the ESS sexual rate. Parameters are: *k*_*f*_ = *k*_*a*_ = 1.2 for panel (1a); *k*_*f*_ = 1.2, *k*_*a*_ = 0.3 for panel (2a); *k*_*f*_ = 1.2, *k*_*a*_ = 0.5for panel (2b); *k*_*f*_ = 0.8, *k*_*a*_ = 1.2 for panel (3a); and *k*_*f*_ = *k*_*a*_ = 0.8 for panel (4a). For all panels, *α* = 0, 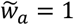, and *k*_*m*_ = 1.5.

For separate-sex populations, we assume that female individuals can reproduce asexually or produce female gametes, whereas males produce only male gametes, and fertilization occurs randomly. Modifier mutants may arise in both sexes. Let *x*_*i*_ (*i* = *AA, Aa, aa*) be the genotype frequency among females, and *y*_*i*_ the frequency among males that successfully sire these females. The recursions of genotype frequencies are then given by

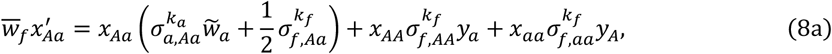

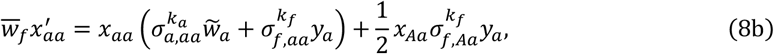

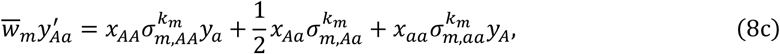

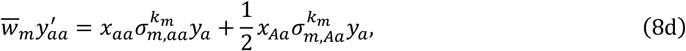

where 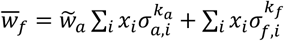 is the mean fitness and 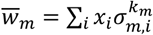 is the mean male fertility. The recursion of allele frequency of the mutant 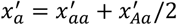 and 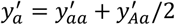. Since rare modifier mutants initially can only be in heterozygous state, with small modification of the sexual rate (|Δ*σ*| ≪ 1), the leading eigenvalue is identical to that for hermaphroditic populations with no selfing (Section 1 of the SI Appendix), and thus the invasion conditions are the same.

To examine the invasion dynamics and relax the assumption of small modification effects, we carried out simulations. Briefly, for a population with initial allocation state *σ*_0_ and *F*_0_, a new mutant arises at each time step. The modification effects of the mutant, Δ*σ* and Δ*F* = *f* − *F*, are drawn from uniform distributions as Δ*σ*/[*σ*(1 − *σ*)] ∼ Unif(−Δ*σ*_max_, Δ*σ*_max_)and Δ*F*/[*F*(1 − *F*)] ∼ Unif(−Δ*F*_max_, Δ*F*_max_), where Δ*σ*_max_ and Δ*F*_max_ denote the maximum magnitudes of modification. Whether a mutant invades is determined by the leading eigenvalue. If a mutant successfully invades, the allocation state of the population is updated to that of the mutant.

## Supplementary Information for

### This PDF file includes

Supporting text

Figure S1

## 1. Invasion analysis

Below I analyze the invasion condition of sex, by distinguishing between whether the residential population is obligately asexual or not. The invasion condition of sex differs under the two scenarios. This is because the sexually reproducing mutants are initially rare in the population, so their male gametes cannot sire residents and female gametes cannot be fertilized when the residents are obligately asexual, unless they can self-fertilize.

### Obligately asexual population (σ = 0)

When *σ* = 0, assuming the mutant initially can arise as heterozygote *Aa*, by Equations (7a) and (7b), the allele frequency change after one generation is

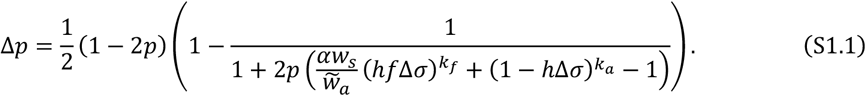

The invasion of sex (i.e., Δ*p* > 0) requires

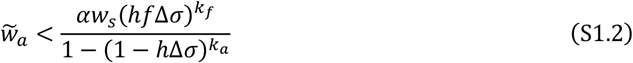

By Equation (S1.2), we can analyze the effects of return exponent on the invasion condition. The effect of the return exponent of female fertility *k*_*f*_ and asexual fertility *k*_*a*_ are, respectively,

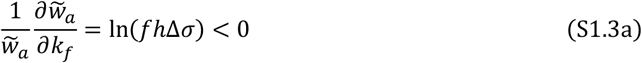

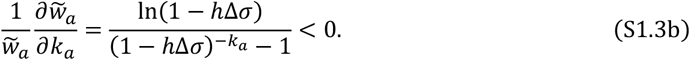

Therefore, higher values of *k*_*f*_ and *k*_*a*_ inhibit the evolution of sex.

### Non-obligately asexual population (σ > 0)

For hermaphroditic populations, assuming small modification of the sexual rate (|Δ*σ*| ≪ 1), the leading eigenvalue of recursion Equations (5a) and (5b), *λ*, is

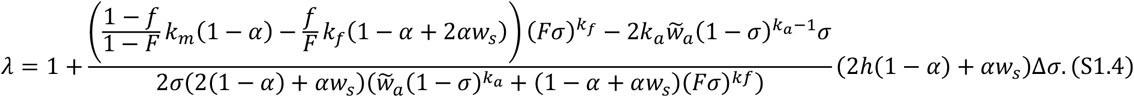

Equation (S1.4) is an exact solution under random mating *α* = 0 even for large Δ*σ*. Solving *λ* − 1 > 0 gives the condition given by Equation (1).

### Separate-sex population

For separate-sex populations, assuming |Δ*σ*| ≪ 1, by Equations (7), the leading eigenvalue of the recursions of the modifier allele frequency within females and males is

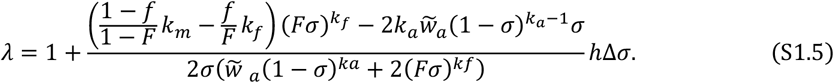

Equation (S1.5) is identical to Equation (S1.4) with *α* = 0.

## 2. Effects of key factors on the invasion of sex when *σ*>0

### Effects of return exponents

By Equation (2), it can be directly seen that a higher return exponent of male fertility, *k*_*m*_, promotes the invasion of sex by increasing the benefit from siring success. For the impact of return exponent of asexual fertility, *k*_*a*_, its effect on the critical fitness *w*_*c*_ is

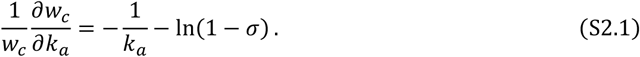

Therefore, a higher *k*_*a*_ promotes the invasion of sex 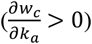 when 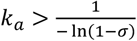. This means that high values of *k*_*a*_ tend to inhibit the invasion of sex when the background sexual rate *σ* is low, while promotes it when *σ* is high.

Assuming modification of sexuality nearly keeps the original sex allocation within sexual reproduction (*f* ≈ *F*), the effect of return exponent of female fertility *k*_*a*_ on *w*_*c*_ is

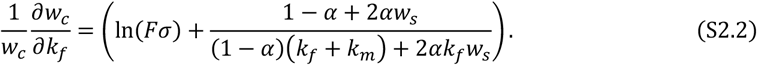

Equation (S2.2) shows that an increase in *k*_*f*_ has opposing effects on the invasion of sex. The first term in the parenthesis, which is negative, shows that an increase in *k*_*f*_ weakens the benefit of sex via increased female fertility. The second term, which is positive, represents that an increase in *k*_*f*_ increases the benefit of sex from higher male siring success. Overall, a higher *k*_*f*_ promotes the invasion of sex 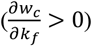 when 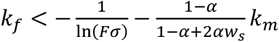, and the right hand side increases with background investment to female function, *Fσ*.

### Effects of background allocation to female function F

We consider two cases, depending on the relationship between *f* and *F*. Assume that *f* and *F* are independent, the effects of *F* on the critical fitness *w*_*c*_ is

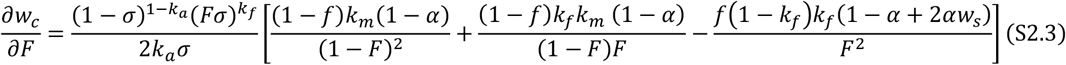

Therefore, the sign of 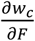 does not depend on the return exponents of asexual fertility *k*_*a*_. When *k*_*f*_ ≥ 1, all three terms in the square bracket are non-negative, so that 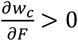, meaning that higher background allocation to female function promotes the invasion of sex (panels (a) and (b) in Figure S1).

A more realistic situation is that modification of sex does not deviate from the original level of sex allocation much (*f* ≈ *F*). In this case, the effect of *F* on the critical fitness *w*_*c*_ is

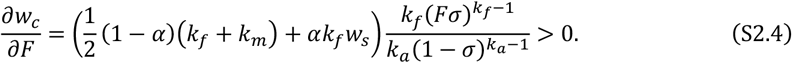

Therefore, a higher background allocation to female function always promotes the invasion of sex.

### Effects of relative modification in female function (f)

The invasion of sex also depends on the relative change in female function within sexual rate modification, *f*. To see how the relative modification to female function, *f*, of a sex modifier affects the invasion of sex, note that the effects of *f* on the critical fitness *w*_*c*_, is

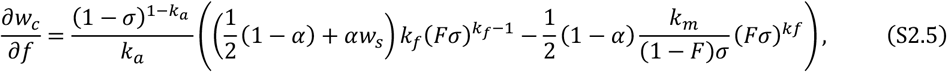

Therefore, 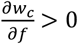 when

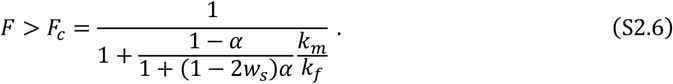

Therefore, provided that the background level of female function allocation, *F*, exceeds a critical level (*F* > *F*_*c*_), the invasion of sex is more likely when modifiers divert more resources to the male function (lower *f*; region to the right of *F*_*c*_ in Figure 2). Conversely, when *F* < *F*_*c*_, a higher *f* promotes the invasion of sex, but in this case, evolution of sex is generally unlikely since it requires the fitness of asexually reproduced offspring to be excessively low (region to the left of *F*_*c*_ in Figure 2). This critical level *F*_*c*_ is lower when the return exponent of male fertility relative to female fertility, *k*_*m*_/*k*_*f*_, is higher. Intuitively, when female fertility of residential individuals is high, the benefit through male siring success outweighs the cost of reduced female fertility.

### Effects of background sexual rate σ

To see the effects of background sexual rate *σ* on the invasion of sex, by Equation (2), we have

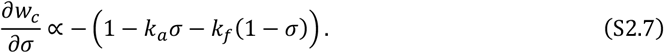

Therefore, when *k*_*a*_ = *k*_*f*_ = 1, the invasion condition of sex is independent of the background sex rate *σ*. When *k*_*a*_, *k*_*f*_ > 1, a higher *σ* always promotes the invasion of sex, and when *k*_*f*_, *k*_*a*_ < 1, a higher *σ* always inhibits the invasion of sex. When *k*_*a*_ > 1 > *k*_*f*_, increasing *σ* from 0, it first inhibits the invasion of sex and then promotes it when 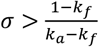. When *k*_*f*_ > 1 > *k*_*a*_, increasing promotes the invasion of sex and then inhibits it when 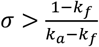.

For biological intuitions, consider a resident population with background sexual rate *σ*. For residential individuals, the fitness obtained via asexual reproduction, female function and male functions is 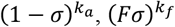 and 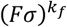 respectively. Consider a sex modifier mutant with effect Δ*σ*. Compared with residents, the loss of fitness via asexual reproductive, and the fitness benefit obtained via female and male functions is respectively,

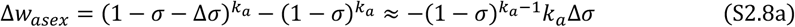

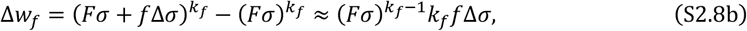

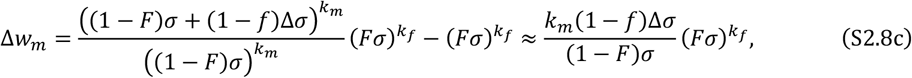

where the approximation assumes small modification (Δ*σ* ≪ 1). In Equation (S2.7c), 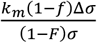 is the increased contribution to the male gamete pool due to increased male gamete production, and 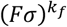 is the female gametes available to be fertilized.

Based on Equation (S2.7), we can analyze the effects of background sexual rate *σ* on the cost and benefit. A special case is that the cost of reduced asexual reproduction is independent of the background sexual rate *σ* when the return exponent is linear for asexual reproduction (*k*_*a*_=1). Similarly, the benefit via both female and male functions is independent of *σ* when the return exponent of female function is linear (*k*_*f*_=1).

By Equation (S2.7a), the effects of background sexual rate *σ* on the cost of reduced asexual reproduction is 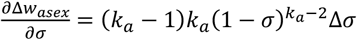. Therefore, when asexual fertility exhibits accelerating returns with investment (*k*_*a*_ > 1), the cost is weaker with higher background sexuality. For the benefit via female function, by Equation (S2.7b), 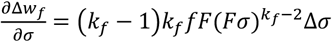, so that a higher background sexual rate *σ* increases the benefit when female fertility shows accelerating returns with investment (*k*_*f*_ > 1), while reduces the benefit when *k*_*f*_ < 1.

To see how the background sexual rate *σ* affects the benefit of sex modifiers via male function, by Equation (S2.7c), we have

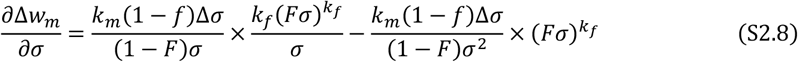

Equation (S2.8) reveals the opposing effects of background sexuality *σ* on the benefit from siring success. The first term means that an increase in *σ* makes more female gametes available to be sired. The second term, which is negative, shows that a higher *σ* means more male gamete production of the resident, which weakens the benefit of a sex modifier via higher contribution to the male gamete pool (i.e., 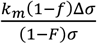). The ratio of the first term to the second term in Equation (S2.8) is *k*_*f*_. When *k*_*f*_ > 1, the positive effect outweighs the negative effect, thus a higher background sexual rate increases the benefit via male function.

To summarize, when *k*_*a*_ = *k*_*f*_ = 1, the benefit of sex modifier from increased female fertility and male siring success is independent of *σ*, so the invasion condition of sex is independent of background sex rate. When *k*_*f*_, *k*_*a*_ > 1, a higher background sex rate *σ* promotes the invasion of sex because it increases the benefit of sex modifier through both female and male functions, and also weakens the cost of reduced asexual fertility. When *k*_*f*_ < 1 and *k*_*a*_ < 1, a higher *σ* inhibits the invasion of sex because it reduces the benefit of sex while increases the cost. When *k*_*f*_ > 1 > *k*_*a*_ or *k*_*f*_ < 1 < *k*_*a*_, an increase in *σ* simultaneously increases or reduces both the benefit and the cost, and whether it promotes the invasion of sex depends on the original sexual rate.

## 3. Evolutionarily stable state (ESS) analysis

Consider a mutant with modification Δ*σ* in a population with state *σ* and *F*. Let’s first consider the case when modification of sexuality does not alter sex allocation (i.e., *f*=*F*). Assuming small modification effect (|Δ*σ*| ≪ 1), an allocation state is evolutionarily stable when 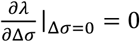 and 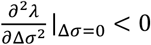 (Maynard Smith & Price 1972), where *λ* is the leading eigenvalue. The first condition holds when *F* and *σ* satisfy the relationship

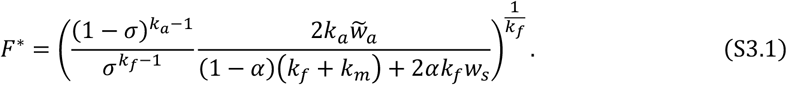

The population with an initial sexual rate *σ* will evolve to lower sexual rate when *F* < *F*^∗^(*σ*) and higher sexuality when *F* > *F*^∗^(*σ*). For the special case when both asexual fertility and female fertility have linear return with investment, *k*_*a*_ = *k*_*f*_ = 1, *F*^∗^ is independent of the sexual rate *σ*.

To examine convergence stability, let 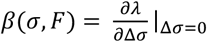 be the selection gradient. The ESS is convergence stable when 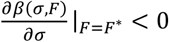. It can be shown that

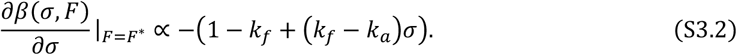

Therefore, when *k*_*f*_, *k*_*a*_ > 1, 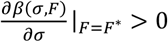 for all 0 ≤ *σ* ≤ 1, so that the ESS line is convergence unstable, and a population initially will evolve to lower sexual rate and finally to *σ* = 0 when *F* < *F*^∗^(*σ*), and to *σ* = 1 when *F* > *F*^∗^(*σ*). Conversely, when *k*_*f*_, *k*_*a*_ < 1, 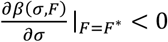, so that the ESS line is convergence stable. When *k*_*f*_ > 1 > *k*_*a*_, the ESS line is convergence stable when 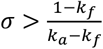, while convergence unstable for the part 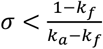. When *k*_*f*_ < 1 < *k*_*a*_, the ESS line is convergence stable for 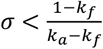 while unstable for 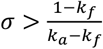.

Now consider the case when the modification of sexual rate may also alter sex allocation (*f* ≠ *F*), and let Δ*F* = *F* − *f*. Note that in our model, modifiers may change sex allocation only when it changes the sexual rate, and there is no mutant that changes sex allocation without changing sexuality, so that 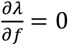. The leading eigenvalue can thus be expanded as 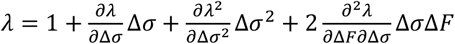, where the derivatives are evaluated at Δ*σ* = Δ*F* = 0. An ESS that is convergence stable exists when 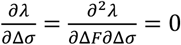 and 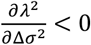. The conditions 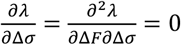 solves to Equations (S2.1) and the ESS sexual rate, given by

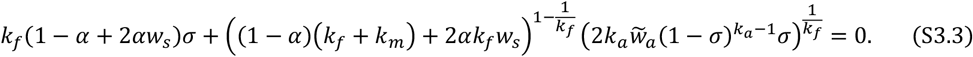

Analytical solution to Equation (S3.3) is available only for special cases. When *k*_*a*_ = 1, Equations (S3.1) and (S3.3) give the ESS state

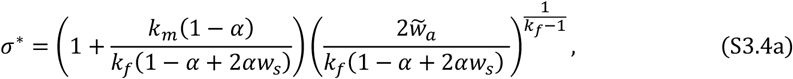

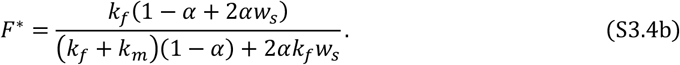

## 4. Effects of selfing on the invasion condition and ESS

By Equation (2), the effect of selfing rate *α* on the threshold fitness, *w*_*c*_, is

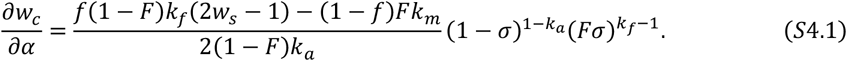

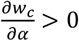 solves to

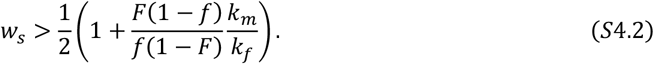

To see how selfing influences the ESS, we need to discuss different cases based on the return exponents of asexual and female fertility. When *k*_*f*_, *k*_*a*_ ≥ 1, populations will evolve to either obligate sexuality or asexuality, depending on whether the initial female function allocation *F* is above or below a crititcal level *F*^∗^, given by Equation (S3.1). By Equation (S3.1), we have

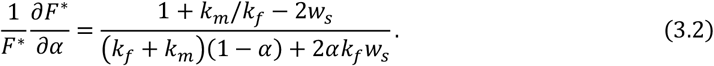

Selfing expands the parameter space where populations evolve to obligate sexuality 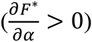 when 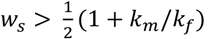. As selfed offspring tend to be less fit than outcrossed offspring, this condition may hold only when male fertility when male fertility exhibits lower returns than female function (*k*_*m*_ < *k*_*f*_), a scenario that may apply to many plant populations (Zhang 2006).

When *k*_*f*_ ≥ 1 and *k*_*a*_ < 1, populations will always evolve to obligate asexuality when the return exponent of asexual fertility, *k*_*a*_, exceeds a critical level, and can evolve to either obligate asexuality or facultative sexuality when *k*_*a*_ is below this level (see case (3) in Table 2). Selfing increases this critical level when the fitness of selfed offspring is high enough (compare red, blue and green lines with black lines in Figure S1a).

When *k*_*f*_ < 1 and *k*_*a*_ ≥ 1, selfing increases the parameter space where populations evolve to facultative sexuality, and makes evolution to obligately sexuality less likely (Figure S1b). In all cases, when the ESS is facultative sexuality, selfing reduces the ESS sexual rate unless *w*_*s*_ is sufficiently high (Figure S1).

## Supplementary Figure

**Figure S1.**
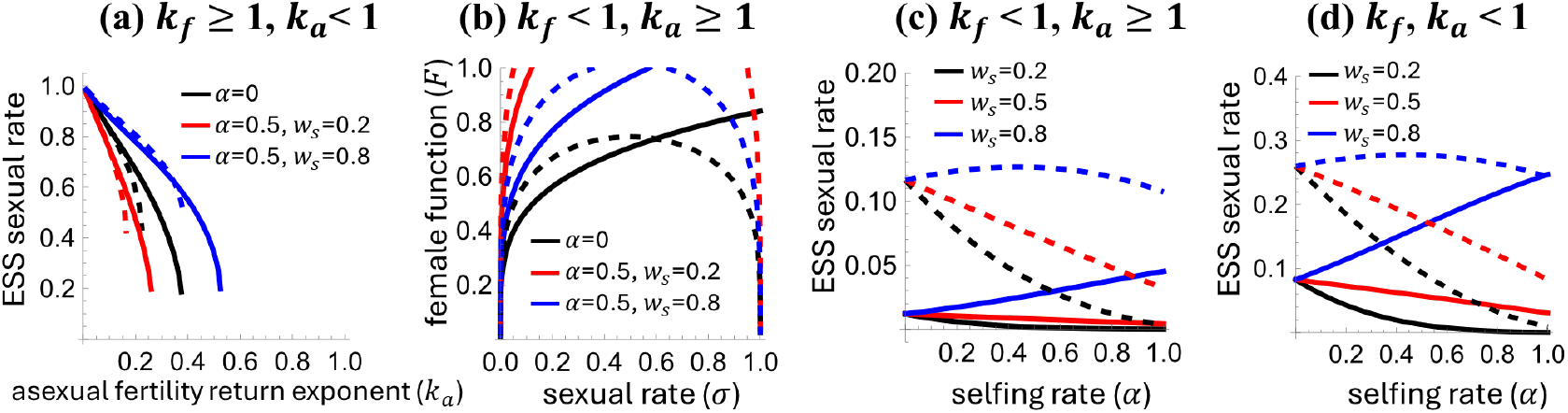
Impacts of self-fertilization on the ESS sexual rate under three cases of the return exponents of female fertility (*k*_*f*_) and asexual fertility (*k*_*a*_) (see Table 2). When *k*_*f*_ ≥ 1, *k*_*a*_< 1, the ESS is obligately asexual when *k*_*a*_ exceeds a critical value, and obligately asexual or facultative sexual when *k*_*a*_ is below this value. Panel (a) shows the influence of selfing on this critical value (x axis at which the curve is truncated) and the ESS sexual rate, where *k*_*f*_=1.1 for solid lines and *k*_*f*_=1.5 for dashed lines. When *k*_*f*_ < 1, *k*_*a*_ ≥ 1, populations evolve to obligately sexual or facultative sexual, depending on the initial state. Panel (b) show the influence of selfing on the parameter space where populations will evolve to the two ESS, where the curve is the critical female function allocation *F*^∗^ truncated below 1, and populations evolve to higher sexual rates when *F* < *F*^∗^, and to lower sexual rates when *F* > *F*^∗^. Panel (c) shows the effects of selfing rate on the ESS sexual rate when facultative sexuality is ESS. In panel (b), *k*_*f*_=0.8, and *k*_*a*_=1 for solid lines and *k*_*a*_=1.2 for dashed lines. In panel (c), *k*_*a*_=1.2, and *k*_*f*_=0.8 for solid lines and *k*_*f*_=0.6 for dashed lines. When *k*_*f*_, *k*_*a*_<1, populations always evolve to facultative sexuality, and panel (d) shows the effects of selfing rate on the ESS sexual rate, where *k*_*a*_ =0.8, and *k*_*f*_=0.8 for solid lines and *k*_*f*_=0.6 for dashed lines. For all panels 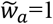, *k*_*m*_=1.5.

